# Integrated proteome and lipidome analyses place OCIAD1 at mitochondria-peroxisome intersection balancing lipid metabolism

**DOI:** 10.1101/2024.11.15.623757

**Authors:** Vanessa Linke, Mateusz Chodkowski, Kacper Kaszuba, Mariusz Radkiewicz, Vikas Rana, Dorota Stadnik, Michał Dadlez, Agnieszka Chacinska

## Abstract

OCIAD1 (Ovarian Cancer Immunoreactive Antigen Domain Containing 1) is a membrane protein largely localized to mitochondria, however, its function in health or disease is not well understood. To comprehensively characterize the molecular changes upon lack of OCIAD1, we used mass spectrometry to study the mitochondrial and cellular proteome and lipidome. We find extensive lipidome rearrangement in OCIAD1 KO cells, characterized by two main phenotypes of decreased ether phospholipids and decreased phospholipids with an odd number of carbons. The lipidomic changes suggest alterations in peroxisomal lipid metabolism. At the same time, proteins responsible for mitochondrial fatty acid β oxidation are significantly increased. Together with a global loss in peroxisomal proteins and a meta-analysis of proximity labeling data, this gives a function to the previously observed partial localization of OCIAD1 to peroxisomes. We suggest a role for OCIAD1 in balancing mitochondrial and peroxisomal lipid metabolism, and a direct impact on the key enzymes FAR1 and ACBD3.

**Summary Statement:** Lipidomics and proteomics of mitochondrial fractions and whole cells lacking the membrane protein OCIAD1 suggest a role as a dually localized protein balancing mitochondrial and peroxisomal lipid metabolism.

## INTRODUCTION

OCIAD1, also known as Asrji, is one of two OCIA domain containing membrane proteins whose function is not well understood. Physiologically, OCIAD1 has been linked to cancer, a connection from which also the domain name Ovarian Carcinoma Immunoreactive Antigen stems (Wang et al. 2010). Early reports in Drosophila have placed OCIAD1 at a variety of organelles along the endocytosis pathway (Kulkarni et al. 2011), however, most human studies have described OCIAD1 to localize primarily to mitochondria, with reports localizing it to the outer membrane (Lee et al. 2017; Antonicka et al. 2020; Elancheliyan *et al*. 2024) and inner membrane (Le Vasseur et al. 2021). Compared to its mitochondrial localization, a smaller pool of OCIAD1 has been localized to peroxisomes in two separate high-throughput studies (Antonicka et al. 2020; Singin et al. 2024). Peroxisomes partly originate from mitochondria, collaborate with mitochondria in lipid metabolism and share a subset of proteins, including for the fission machinery (Costello et al. 2018). OCIAD1 has been described as modulating mitochondrial and peroxisomal morphology by limiting mitochondrial network formation and peroxisome numbers (Antonicka et al. 2020; Singin et al. 2024).

OCIAD1 contains two transmembrane domains and an extended C-terminal tail that has been found to interact with prohibitins (Yoshinaka et al. 2019; Le Vasseur et al. 2021; Elancheliyan *et al*. 2024). Prohibitins are large membrane protein complexes that manage local lipid composition (Lourenço et al. 2021). Further and consistent with its transmembrane segment, OCIAD1 has been described in a global study to directly interact with lipids (Niphakis et al. 2015). Despite these indications of an involvement of OCIAD1 in lipid metabolism, lipidomics of cells with altered OCIAD1 levels have so far only been performed once, as part of a high-throughput CRISPR knockout study in whole cell lysates of HAP1 cells (Rensvold et al. 2022).

Very recently, our group introduced and biochemically characterized a knockout of OCIAD1 in HEK 293 cells (Elancheliyan *et al*. 2024). Here, we use this OCIAD1 KO to study the effects of a loss of OCIAD1 on the proteome and lipidome of whole cells and of fractions enriched in mitochondria after challenging the cells by culturing in galactose medium in the absence of glucose (de Kok et al. 2021). We find extensive proteome and lipidome rearrangement, suggesting a role for OCIAD1 in lipid metabolism of mitochondria and peroxisomes.

## RESULTS AND DISCUSSION

### Lipidome rearrangement upon OCIAD1 KO reveals that lack of OCIAD1 affects peroxisomal lipid metabolism

Utilizing a discovery lipidomics approach, we identified 866 lipid species from 30 lipid classes and further quantified an additional 3201 chromatographic features. Upon OCIAD1 KO, 25% of identified lipids (215) are significantly increased in their abundances compared to wildtype cell lysates while 18% (153) are decreased. In the mitochondrial fraction upon OCIAD1 KO, 14% of lipids (120) are significantly increased and 22% (189) decreased. For the mitochondrial isolates, we observed an increase in individual lipid species including the sulfatide SHexCer d36:2, the phospholipid PE 38:2, as well as the glycerolipid Alkenyl-DG P-18:0_20:3, while we observed the biggest decreases in phospholipids including Plasmanyl-PC O-35:1 and Plasmenyl-PC P-33:0 (**Figure 1A**). When analyzing the lipid changes in the mitochondrial fraction upon OCIAD1 KO by lipid class, we saw the biggest decrease in ether-linked Plasmanyl-PC (average FC -1.1) and vinyl ether-linked Plasmenyl-PC (average FC -0.6) species, while the ester-linked PE (average FC +0.3) and PC (average FC +0.2) phospholipids exhibited the biggest increase upon OCIAD1 KO (**Figure 1B**). Most other measured lipid classes, including the mitochondrial lipid cardiolipin (CL), were not markedly changed on average upon OCIAD1 KO.

**Figure 1.**
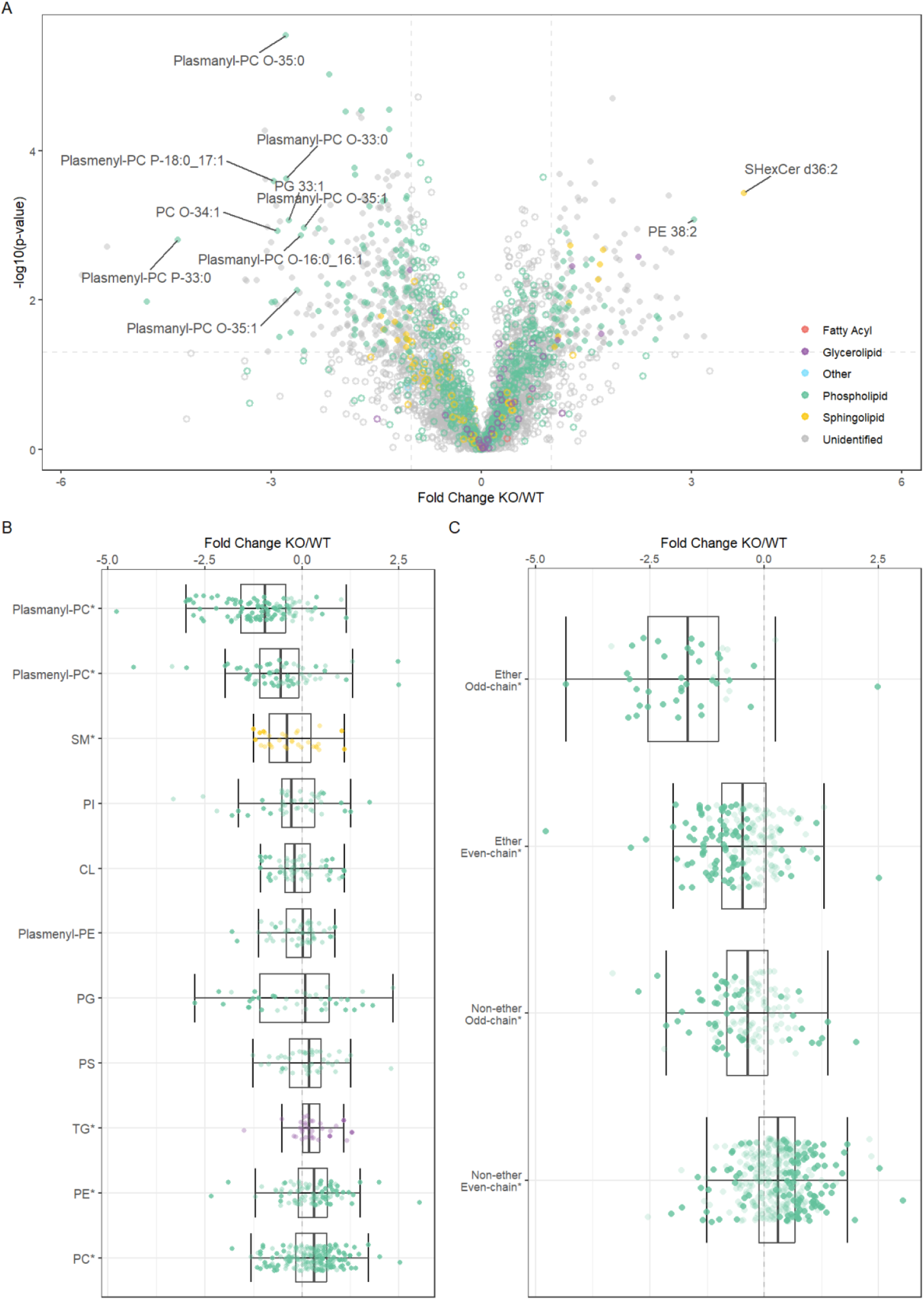
Lipidome changes in the mitochondrial fractions upon OCIAD1 KO show a decrease in ether phospholipids containing an odd number of carbons. Lipidomics of OCIAD1 KO mitochondrial fractions compared to WT represented as (A) volcano plot, (B) boxplot by headgroup (n > 10), ordered by mean FC, and (C) boxplot of ether phospholipids and phospholipids containing an odd number of carbons. Colors refer to lipid class (see legend), asterisks indicate p < 0.05 when comparing to FC = 0, and lower transparency indicates p < 0.05 when comparing OCIAD1 KO to WT.

When further summarizing the phospholipid changes in the mitochondrial fraction upon OCIAD1 KO by fatty acid chain, we generally observed a significant decrease (average FC - 0.7) in ether-linked phospholipids (ePL) (**Figure 1C**). Ether lipids are a major (∼20%) structural component of eukaryotic cell membranes (just like OCIAD1, they are absent in fungi) that alter membrane dynamics towards more rigidity by allowing for tighter packing of phospholipids in the membrane (Dean and Lodhi 2018). Ether phospholipids are synthesized in the peroxisome and ER, thus their decrease upon OCIAD1 KO could indicate a reduced synthesis in peroxisomes and/or the ER . Consistent with a partial localization of OCIAD1 to the peroxisome, it is reasonable to hypothesize a potential direct role of OCIAD1 in supporting peroxisomal ether phospholipid synthesis.

Structurally, ePLs consist of a PC or PE headgroup in the sn3 position and a fatty acid in the sn1 position that is bound via a characteristic ether (“plasmanyl-”) or vinyl ether bond (“plasmenyl-” aka plasmalogens). In the sn2 position, a fatty acid is connected via an ester bond, which classically in ePLs is preferably a polyunsaturated fatty acid (PUFA), commonly 22:6 or 20:4. The largest decrease in ePLs observed in OCIAD1 KO cells, however, is not a decrease in ePLs with PUFAs (average FC -0.4) but rather in ePLs containing an odd number of carbons in the sn2 chain (average FC -1.7), e. g. a decrease in Plasmenyl-PC P-18:0_17:1 (**Figure 1C**). Looking beyond ePLs, we observed a general decrease in phospholipids containing an odd number of carbons (average FC -0.7) that was similar in size to the decrease in ePLs, thus we concluded that decreases in ePLs and decreases in odd chain fatty acids upon OCIAD1 KO were due to additive effects. An odd number of carbons may indicate the presence of - for the mammalian system rare - odd chain fatty acids. However, with the lipidomics method applied here, these are not distinguished from branched fatty acids (BCFAs). The additional methyl group in BCFAs prevents them from undergoing β-oxidation in mitochondria, and instead they are targeted to peroxisomes for α-oxidation (Lodhi and Semenkovich 2014). A decrease in BCFA-PLs upon OCIAD1 KO could thus indicate increased α-oxidation in peroxisomes.

To summarize, we have found multiple links of OCIAD1 to peroxisomal lipid metabolism in the mitochondrial lipidome changes upon OCIAD1 KO. Importantly, these changes were conserved in the corresponding whole cell lysates (Figure S1). Since ether phospholipids are synthesized in peroxisomes, their decrease upon OCIAD1 KO may indicate that OCIAD1 is necessary for peroxisomal ether phospholipid synthesis. It is known that ePLs originating from the peroxisome affect mitochondria in various ways, for example by facilitating supercomplex and respirasome assembly (Bennett et al. 2021). It is at present unknown whether OCIAD1 could regulate these or other assemblies but a function for OCIAD1 in regulating the available lipid composition is conceivable based on our data. On the other hand, the loss of OCIAD1 decreased phospholipids containing an odd number of carbons, possibly BCFA-PLs, which might indicate increased peroxisomal α-oxidation. Together, the lipidomics data upon OCIAD1 KO suggest a role for OCIAD1 in balancing peroxisomal lipid anabolism and catabolism and/or a role for OCIAD1 in lipid transport in and out of peroxisomes.

### Proteins of mitochondrial fatty acid β-oxidation are increased upon OCIAD1 KO

In our bottom-up proteomics analysis of the same OCIAD1 KO cells and their mitochondrial isolates, we identified 3,152 protein groups, of which 741 are known mitochondrial proteins (representing 65% of proteins listed in MitoCarta 3.0 (Rath et al. 2021)). In the mitochondrial fraction, OCIAD1 was barely detectable upon its knockout and certain CHCH-domain containing proteins showed differential regulation, with CHCHD2 being decreased and CHCHD7 increased upon OCIAD1 KO. Overall and in accordance with the recently-described role of OCIAD1 in stress (Elancheliyan et al. 2024), however, the majority of the mitochondrial proteome remained unchanged upon OCIAD1 KO (**Figure 2A**).

**Figure 2.**
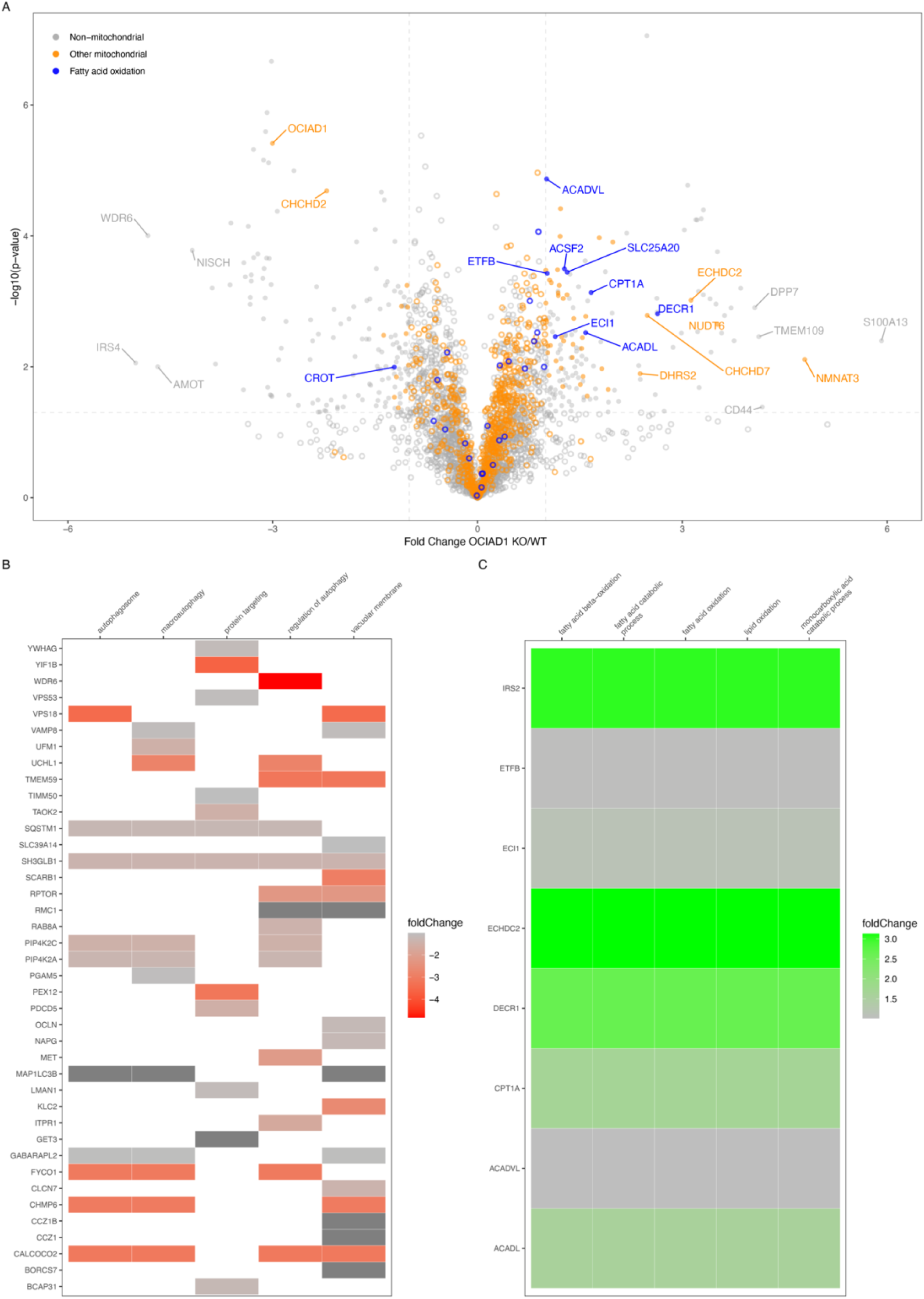
Proteome changes in the mitochondrial fractions upon OCIAD1 KO show an increase in mitochondrial fatty acid β oxidation. Proteomics of OCIAD1 KO mitochondrial fractions compared to WT represented as volcano plot, colored by proteins of mitochondrial fatty acid β oxidation pathway (blue) and other mitochondrial proteins (orange, MitoCarta 3.0) **(A)**. Heatmap of the top 5 enriched GO terms of proteins significantly decreased **(B)** and increased **(C)** upon OCIAD1 KO.

When performing GO term enrichment analysis on the significantly regulated proteins in the mitochondrial fraction upon OCIAD1 KO, lipid metabolic processes, specifically mitochondrial fatty acid β-oxidation, appeared as significantly increased (**Figure 2B**). This phenotype might be related to the lost interaction of OCIAD1 with prohibitins, who are known to impact fatty acid oxidation (Wu et al. 2020). One of the most increased proteins upon OCIAD1-KO was mitochondrial 2,4-dienoyl-CoA reductase (DECR1), catalyzing the rate-limiting step for PUFAs before β-oxidation (Nassar et al. 2020). Decreased GO terms in the mitochondrial fraction upon OCIAD1 KO interestingly featured mostly non-mitochondrial proteins. For the whole cell lysates, decreased proteins were enriched in GO terms related to translation and RNA processing, while increased proteins were enriched in GO terms related to exocytotic vesicles (**Figure S2B**).

### The rate-limiting enzyme of peroxisomal ether phospholipid synthesis is decreased upon OCIAD1 KO

Encouraged by the observations of a defect in peroxisomal lipid metabolism suggested by the lipidomics results, we asked whether there were differences in peroxisomal proteins in the mitochondrial fractions upon OCIAD1 KO. Apart from a distinct decrease in PEX12 and in line with the observed decrease in ePLs, we observed a decrease in proteins responsible for peroxisomal ether phospholipid synthesis, specifically a significant decrease in the rate-limiting enzyme FAR1 (**Figure 3A**). FAR1 is a tail-anchored membrane protein whose levels are regulated by degradation, the mechanism of which has remained a mystery (Honsho et al. 2010; Honsho et al. 2013). Interestingly, very recently, a role has been proposed for OCIAD1 in the degradation of a mitochondrial membrane protein, i. e. TIMM17A by YME1L1 (Elancheliyan *et al*. 2024). A precise mechanism through which OCIAD1 might protect peroxisomal FAR1 from degradation to allow for ePL synthesis and whether it might be similar to its role in mitochondria is at present unknown, however, OCIAD1 is an intriguing new target to study the post-translational regulation of FAR1 levels (Ferdinandusse et al. 2021; Cui et al. 2021; Honsho and Fujiki 2023).

**Figure 3.**
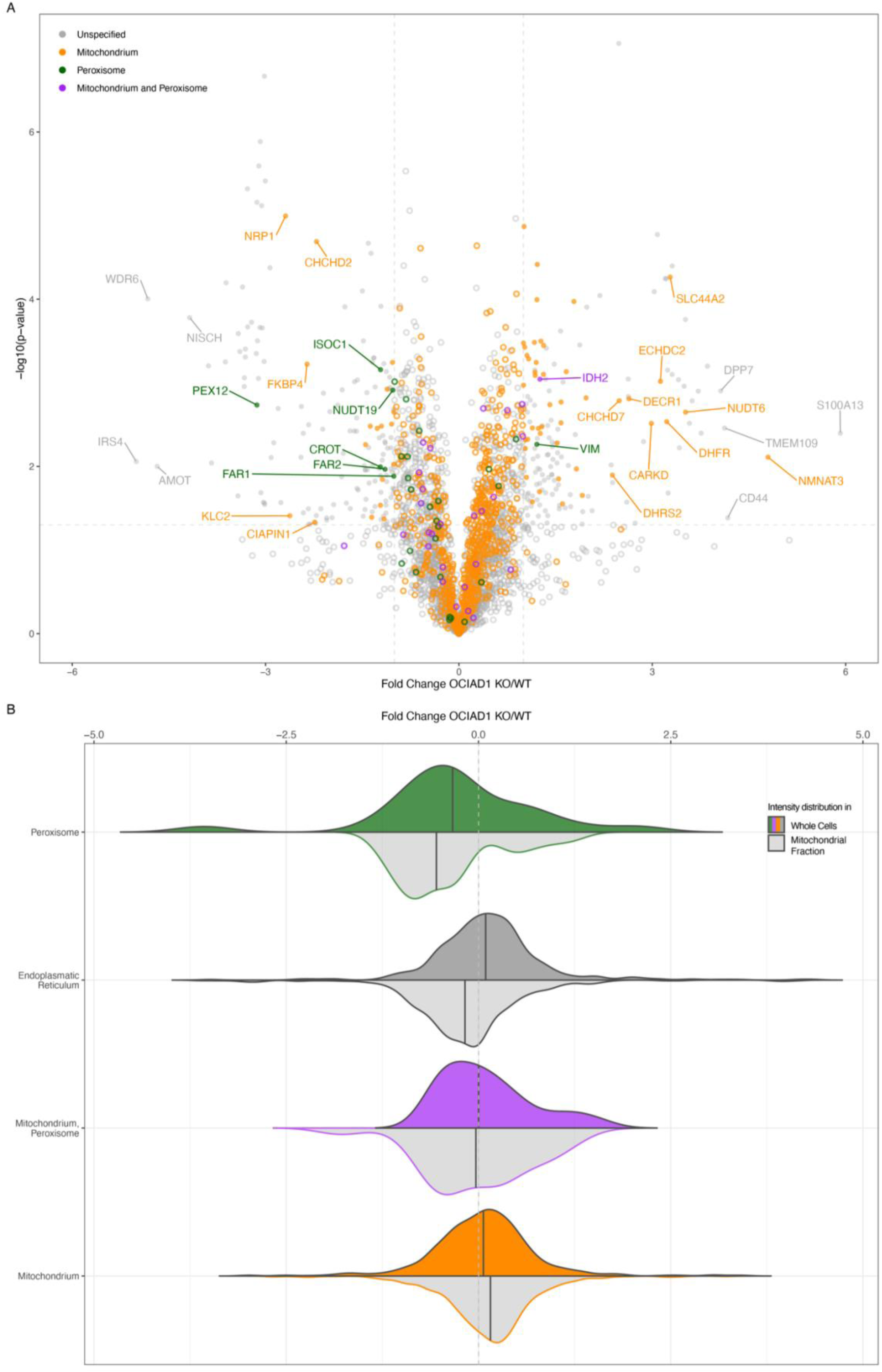
Proteome changes upon OCIAD1 KO by organelle show a decrease in peroxisomal proteins. **(A)** Volcano plot of OCIAD1 KO mitochondrial fractions compared to WT as in Figure 2A but colored by organelle shows a decrease in peroxisomal proteins (green) but not proteins dually localized to mitochondria and peroxisomes (purple). (**B**) When comparing FCs of organellar proteins by sample type, the decrease in peroxisomal proteins is most evident in the mitochondrial fraction.

### Comparative proteome changes by organelle suggest a weakened contact between mitochondria and peroxisomes upon lack of OCIAD1

Other changes concerning peroxisomal lipid metabolism were smaller and below significance but trending down, including enzymes responsible for peroxisomal branched chain fatty acid α-oxidation (below significance decreases in HACL1, ALDH3A2, SLC27A2). This trend was in contrast to the lipidomics results and let us conclude that the decrease in phospholipids with an odd number of carbons upon OCIAD1 KO was not driven by an increase in peroxisomal α-oxidation of BCFAs. However, since these proteomic changes were observed in the mitochondrial isolates of OCIAD1 KO cells, they may not indicate functionally relevant decreases but rather an overall decrease in peroxisome abundance or a decreased contact between peroxisomes and mitochondria. Supporting this hypothesis, only two peroxisomal proteins were found to be significantly increased in the mitochondrial fraction, one of which is dually localized to mitochondria (IDH2), the other filamentous (VIM). Expanding this organellar analysis more generally, in the mitochondrial fraction, we observed a decrease in proteins from peroxisomes that was not the case for proteins from mitochondria, or dually localized proteins (**Figure 3B**). This specific decrease in peroxisomal proteins appeared less strong in whole cell lysates. Further, we saw a similar behavior for ER-resident proteins, albeit with a smaller fold change. In summary, our proteomics data indicates that there might be a small decrease in peroxisomal proteins in OCIAD1 KO cells overall, however, there is a substantial decrease in peroxisomal, as well as to a lesser extent ER proteins, in the mitochondrial isolates of these cells. Together, this led us to hypothesize that there may be a decreased contact between mitochondria and other organelles upon OCIAD1 KO.

### A meta-analysis of proximity labeling studies places OCIAD1 at the mitochondria-peroxisome interface

To further investigate this hypothesis, we turned to two published large scale proximity labeling datasets that included OCIAD1 either as bait or prey (Antonicka et al. 2020; Go et al. 2021). The analysis of the first study found OCIAD1 as a bait interacting with several peroxisomal membrane proteins (PEX13, PEX11B, PEX3, PEX6, PEX1, ACBD5, PEX14, PEX19, ALDH3A2) and with proteins dually localized to peroxisomes and mitochondria (ABCD3, DNM1L, MAVS, USP30, MFF, ATAD1, TMEM135, PEX5) (**Figure 4A-B**, (Antonicka et al. 2020)). Further, when correlating the high-throughput profile of OCIAD1 with the profiles of all other baits, a cluster enriched in mitochondrial outer membrane proteins involved in peroxisomal biology emerged (MTCH1, MARCH5, SLC25A46, MTCH2, RHOT2, FIS1, MAVS, FKBP8, PTPN1) (**Figure 4C**, (Antonicka et al. 2020)). Within this cluster, OCIAD1 showed strongest specificity for ABCD3 and to a lesser extent for PEX14 (**Figure 4D**, (Antonicka et al. 2020)).

**Figure 4.**
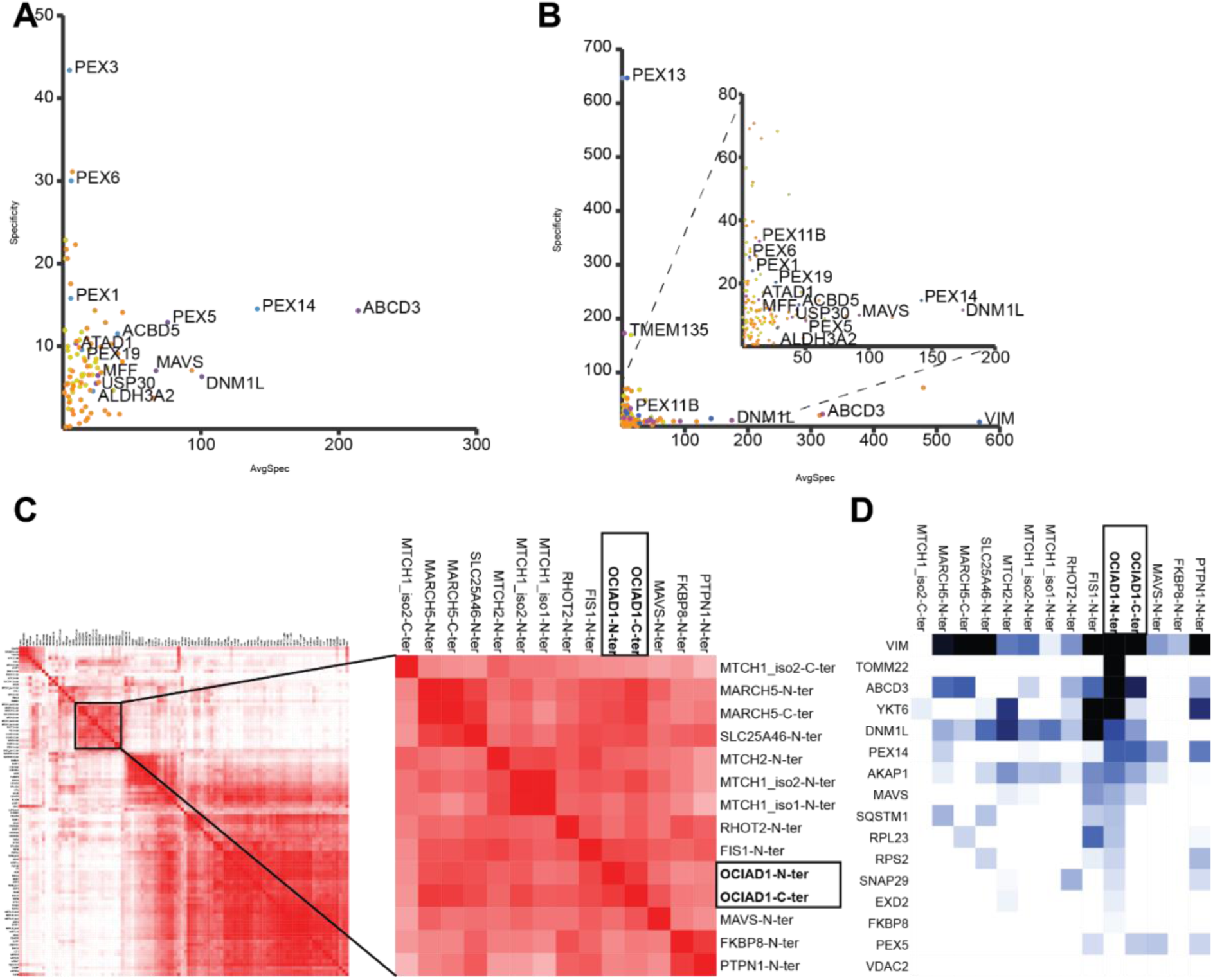
Meta-analysis of proximity labeling data for OCIAD1. Specificity plot of C-terminally tagged OCIAD1 (**A**) and N-terminally tagged OCIAD1 (**B**) as bait, respectively, with peroxisomal proteins (blue), mitochondrial proteins (orange) and dually localized proteins (purple). Baits (**C**) and preys (**D**) correlated with OCIAD1 as bait. Data from Cell Metabolism, Vol.32 (3), Antonicka H, Lin Z-Y, Janer A, Aaltonen MJ, Weraarpachai W, Gingras A-C, Shoubridge EA, A High-Density Human Mitochondrial Proximity Interaction Network, 479– 497.e9, Copyright Elsevier (2020).

The second study included OCIAD1 only as a prey but it showed partially shared interactions with ABCD3 in its proximity to ACBD5, PXMP2 and others (Go et al. 2021). Also in this study, other preys that correlated highly with the behavior of OCIAD1 as a prey included outer mitochondrial membrane proteins (TOMM70, TOMM22, FBXL4, HK1, CISD1, RMDN2, MTFR1, MTFR2, TDRKH, EXD2), peroxisomal membrane proteins (PEX11B, PEX3) and dually localized proteins (USP30, MAVS, DNM1L, RHOT2, MFF, ABCD3, TMEM135, MARC2) (Go et al. 2021).

Together, these proximity labeling studies place OCIAD1 at the outer mitochondrial membrane and peroxisomes and support a possible placement of OCIAD1 at mitochondria-peroxisomal contacts. Emerging as the biggest hit from this meta-analysis, we hypothesize a direct interaction of OCIAD1 with ABCD3, a dually-localized protein responsible for importing specific fatty acid CoAs, including the import of BCFAs CoAs to peroxisomes. This hypothesis is intriguing as it could explain the reduction in phospholipids, specifically peroxisomal ePLs, containing an odd number of carbons upon OCIAD1 KO. We thus propose that, rather than affecting peroxisomal α-oxidation, OCIAD1 may be needed for the import of BCFAs CoAs into peroxisomes via ABCD3.

Besides fission from existing peroxisomes, new peroxisomes can also be formed *de novo* (Banerjee and Prinz 2023). In humans, peroxisomes can be built from PEX16-containing vesicles from the ER and PEX3- and PEX14-containing vesicles from mitochondria, however, open questions remain (Sugiura et al. 2017). Curiously in this regard is the very recently described function of MARCH5 in the formation of pre-peroxisomes from mitochondria (Zheng et al. 2022; 2024). Not only did MARCH5 correlate highly with OCIAD1 in our proximity meta-analysis, in fact, OCIAD1 was found to be a significant interactor of MARCH5 alongside ABCD3, PEX19 and PEX3 (Zheng et al. 2022). Together with the observed decrease in the E3 RING-finger complex member PEX12 upon OCIAD1 KO that may imply reduced peroxisomal matrix import (Rudowitz and Erdmann 2023), we thus hypothesize a role for OCIAD1 in peroxisome biogenesis from preperoxisomal vesicles.

Organellar contacts are commonly established to facilitate protein or lipid trafficking between organelles and are especially vital for peroxisomes (Shai et al. 2016). Dually localized proteins, a group to which OCIAD1 shall be counted, are an important contributor to organellar tethers (Freitag et al. 2024). Curiously, the second high-throughput study used for our meta-analysis suggested OCIAD1 to be localized at the mitochondria-ER interface alongside other outer membrane proteins such as AKAP1, MAVS, and HK1 (Go et al. 2021). ER-peroxisome contact is known to be established via VAPA/VAPB and ACBD4/ACBD5 (Kors et al. 2022), the latter of which was also found proximal to OCIAD1. A three-way interaction between peroxisomes, mitochondria, and the ER has been suggested (Costello et al. 2018), and it is tempting to speculate that the characteristic extended C-terminal arm of OCIAD1 allows for far-reaching interactions supporting a role at the interface of organelles.

### Conclusions

We showed a decrease in ether phospholipids upon OCIAD1 KO supported by proteomics measurements that showed a decrease in the rate-limiting enzyme for peroxisomal ether lipid biosynthesis, FAR1. We also showed a decrease in phospholipids containing an odd number of carbons, likely branched chain fatty acids, that might be less imported into peroxisomes upon loss of a possible interaction of OCIAD1 with the transporter ABCD3. Overall, we observed a decrease in peroxisomal proteins, pointing to a partial loss in contact to mitochondria. In contrast, we saw an increase in mitochondrial fatty acid β oxidation. In this context, it is relevant to consider other species, such as fungi, which do not contain a protein similar to OCIAD1, and neither do they contain ether phospholipids. While in humans, fatty acid oxidation is split between mitochondria and peroxisomes, in fungi and plants it is confined to peroxisomes. With gaining a multi-localized fatty acid oxidation, organisms also needed to develop a way to control and balance these processes. Further supported by a meta-analysis of the behavior of OCIAD1 upon proximity-labeling, our data pinpoints to a role of OCIAD1 in mitochondria-peroxisome contact sites and in the biogenesis of peroxisomes. In summary, this role of OCIAD1 as a key protein in balancing organellar lipid metabolism at the intersection of mitochondria and peroxisomes opens up new exciting avenues for further research which is particularly important to study as it may shed light on the roles of OCIAD1 in human diseases.

## MATERIALS AND METHODS

### Cell culture

Approximately 22.5 million Human Embryonic Kidney (HEK 293, ATCC) wildtype (WT) or OCIAD1 KO cells (n = 3 replicates each, see Elancheliyan *et al*. 2024 for details to OCIAD1 KO cells) were seeded and cultured for one day in Dulbecco’s Modified Eagle’s Medium (DMEM), followed by one day in low glucose DMEM (5.55 mM glucose), followed by one day in galactose-containing DMEM (10 mM galactose, glucose-free) before harvest. All media were supplemented with 10% fetal bovine serum, 2 mM glutamine and 1:100 penicillin-streptomycin. For harvest, cells were taken from 37°C and media was removed. Cells were washed with PBS twice, followed by centrifugation for 10 min at 1000 g at room temperature. Whole cell samples were obtained by preserving 1/10th of each cell pellet.

### Isolation of mitochondrial fraction

Fractions enriched in mitochondria were obtained from the remaining cell pellets using a limited swelling approach as reported earlier (Panov 2013, Elancheliyan *et al*. 2024). All steps were performed on ice. Briefly, the remaining pellet was resuspended in isotonic buffer with 2mM PMSF (75 mM Mannitol, 225 mM Sucrose, 10 mM MOPS, pH 7.2, 1 mM EGTA). After a second centrifugation at 1000 g for 5 min, 4°C, the pellet mass was calculated to resuspend pellets in 5ml hypotonic buffer with 2 mM PMSF (100 mM Sucrose, 10 mM MOPS, pH 7.2, 1 mM EGTA) per g cell pellet. The mixture was incubated on ice for ∼6 minutes and then gently homogenized by 10 strokes in a glass-glass Dounce homogenizer. Then, hypertonic buffer (1.25 M Sucrose, 10 mM MOPS, pH 7.2) was added at a ratio of 1.1 ml per gram cell pellet and the solution gently mixed. Lastly, the volume was tripled with isotonic buffer with 2mM PMSF. The suspension was centrifuged for 10 min at 1000 g, 4°C. To obtain a fraction enriched in mitochondria, the supernatant was centrifuged at 10,000 g for 10 min, 4°C. The pellet was then washed with isotonic buffer and again centrifuged at 10,000 g for 10min/4°C. The protein concentration was determined using the Bradford assay, after which equal amounts were aliquoted and pellets flash frozen in liquid nitrogen and stored at -80°C until analysis.

### Lipidomics sample preparation and LC-MS analysis

Lipid extraction from the collected cell pellets or mitochondrial fractions was performed as previously published (Matyash et al. 2008; Linke et al. 2020). Briefly, samples were thawed on ice and underwent protein precipitation by adding 225 µL of cold methanol and vortexed for 1 min. Next, 750 µL of cold methyl tert-butyl ether (MTBE) was added for lipid extraction and 188 µL of cold water added to induce phase separation. Samples were vortexed for 10 min at 4°C using a ThermoMixer (Eppendorf). Lastly, the samples were centrifuged at 6500 g at 4°C. 700 µL of the upper layer was transferred into a new 1.5 mL sample tube, evaporated to dryness under nitrogen stream at 50°C. The remaining sample was kept for further proteomics analysis.

The dried residue was reconstituted in 100 µL of methanol/toluene solution (9:1, v/v). LC-MS lipidomic analysis was performed using Acquity UPLC system (Waters) coupled with a Q Exactive Orbitrap mass spectrometer (Thermo Scientific). Chromatographic separation was achieved using an ACQUITY UPLC BEH C18 1.7 µm, 2.1 mm × 100 mm column (Waters). Mobile phase A consisted of ACN/H2O (7:3, v/v) with 10mM ammonium acetate and mobile phase B consisted of IPA/ACN (9:1, v/v) with 10mM ammonium acetate. The chromatographic gradient started at 2% B for 2 min, next changed to 30% B to 5 min and up to 50% to 6 min. Percentage of B increased up to 85% to 20 min and up to 95% to 21 min, and held till 28 min. Column was re-equilibrated for the next 7 min. The flow was maintained at 0.4 mL/min and the column oven was set to 45°C.

The mass spectrometer was operated either in positive and negative electrospray ionization mode (ESI). The HESI source was kept at 300°C, sheath gas was set to 25, auxiliary gas flow to 10 and sweep gas flow to 5. The capillary temperature was held at 320°C and the S-lens RF level was set to 55. Capillary voltage in ESI+ was set to 3.4 kV and in ESI-to 4.0 kV. Mass spectra were acquired using data-dependent acquisition (DDA): MS1 resolution was 70,000, MS2 resolution was 35,000. Automatic gain control (AGC) was set to 1.0×10^6^ for MS1 and 5.0×10^5^ for MS2. Maximum injection time was 120 ms for MS1 spectra and 80 ms for MS2 spectra. Dynamic exclusion was set to 10 s with a minimum AGC target at 8.0×10^3^. Ion fragmentation was achieved using stepped normalized collision energy (NCE) set to 20, 30 and 45 with an isolation window of 1.5 m/z.

### Lipidomics data analysis

Lipidomics data analysis was performed in accordance with previously published workflows with several modifications.(Hutchins et al. 2018) Compound Discoverer 3.3.3 (CD, Thermo Scientific) was used for the analysis of mass spectra, chromatographic peak alignment and quantification of the area-under-curve (AUC) for individual features. Two separate methods were run to result in “Aligned” and “Unaligned” features set. For the “Aligned” data, sequence of algorithm workflow was: *Input Files → Select Spectra → Align Retention Times → Detect Compounds → Group Compounds → Fill Gaps → Mark Background*. In the “Select Spectra” *MS(n-1) Precursor* was selected to use the direct parent scan of the spectrum. Retention time alignment in the “Align Retention Times” *Adaptive curve with Linear Model* and *Maximum Shift* of 0.6 min were selected, with *Mass Tolerance* at 7.5 ppm. For the compound detection in “Detect Compounds” *Mass Tolerance* was set to 7.5 ppm, *Minimum Peak Intensity* threshold to 10,000, *Precursor Mass Tolerance* 0.025 Da for [M+H]+1 and [M-H+TFA]-1 precursor ions. Next, grouping of detected compounds was achieved by using “Group Compounds” with *Mass Tolerance* set to 7.5 ppm, *RT Tolerance* to 0.2 min, *Minimum Valley* between the two distinct compounds to 10% for *Preferred Ions* of [M+H]+1 and [M-H+TFA]-1. *Peak Rating Threshold* was set to 0. “Fill Gaps” operated with 7.5 ppm mass tolerance and *S/N Threshold* of 1.5. The *Maximum Sample/Blank* ratio was set to 3 in “Mark Background”. For the “Unaligned” data, sequence of *Input Files → Select Spectra → Detect Compounds* was performed. The analysis parameters were identical to the respective algorithms of the “Aligned” workflow. “Aligned” Compound Discoverer data was exported as excel files using three levels of *Compounds → Compound per File → Features*, and only *Features* for “Unaligned” data.

For lipid identification and quantitation Lipidex 1.1 was employed(Hutchins et al. 2018). RAW files with mass spectra were converted to MGF files using MSConvertGUI (ProteoWizard, Stanford University). They were next uploaded to the *Spectrum Searcher* of Lipidex. Lipid search was performed using in silico built-in *Lipidex_HCD_Acetate* and in-house *Ganglioside* databases. Finally, the “Aligned” and “Unaligned” data sheets from CD were selected for *Peak Finder* algorithm. *Minimum Lipid Spectra Purity* was set to 75%, *Min. MS2 Search Dot Product* to 500, *Min. MS2 Search Rev. Dot Product* to 700m with *FWHM Window Multiplier* set to 2.0 and *Max. Mass Difference* to 15 ppm. In the *Result Filtering Parameters* both *Adduct/Dimer Filtering* and *In-source Fragment Filtering* were enabled with the *Max. RT M.A.D Factor* set to 3.5.

### Proteomics sample preparation and analysis

500 µL MeOH were added to each sample after lipidomics extraction. Samples were then centrifuged at 14 000g for 10 min at 4C and the supernatant removed. After 100 µl lysis buffer (8 M Urea, 100 mM Tris pH 8.5) was added to the pellet and the mixture sonicated, the samples were centrifuged at max. speed and the supernatant diluted 5 times with 100 mM HEPES, pH 8.0 containing 10 mM TCEP and 15 mM CAA. Proteins were digested overnight at 37°C with trypsin (Promega V5113). Then, samples were acidified with TFA to a final pH of ∼ 2 and peptides were desalted using STAGE tips of three layers AttractSPE Discs Bio C18 (Affinisep) (Myers et al. 2019). The resin was conditioned with methanol and equilibrated with 150 μl of 0.5% formic acid (FA). Approximately 30 μg of digested peptides were loaded and washed with 150 μl of 0.1% FA followed by elution with 60 μl of 60% ACN in 0.1% FA. The solvents were removed by SpeedVac after which peptides were dissolved in 2% of acetonitrile in 0.1% TFA before LC-MS/MS measurement.

Chromatographic separation was performed using an Easy-Spray Acclaim PepMap column (50 cm long × 75 μm inner diameter, Thermo Fisher Scientific) at 45 °C by applying 120 min acetonitrile gradients in 0.1% aqueous FA at a flow rate of 300 nl/min. An UltiMate 3000 nano-LC system was coupled to a Q Exactive HF-X mass spectrometer via an easy-spray source (Thermo Fisher Scientific). The mass spectrometer was operated in data-dependent acquisition mode with survey scans acquiring at a resolution of 120,000 at m/z 200. Up to 15 of the most abundant isotope patterns with charge states <6 from the survey scan were selected with an isolation window of 1.3 m/z and fragmented by higher-energy collision dissociation (HCD) with normalized collision energies of 27. Dynamic exclusion was set to 35 s, and maximum ion injection times for the survey and MS/MS scans (acquired with a resolution of 15,000 at m/z 200) were 45 and 28 ms, respectively. The AGC target for MS was set to 3e6 and for MS/MS to 1e5, and the intensity threshold for MS/MS was set to 1.6e4.

Data was processed with MaxQuant v. 2.0.3.0, and the peptides were identified from the MS/MS spectra searched against the UniProt KB Human Proteome (downloaded on 18.09.2023) using the built-in Andromeda search engine. Cysteine carbamidomethylation was set as a fixed modification, and methionine oxidation and N-terminal acetylation were set as variable modifications. Label free quantification was performed and match between runs was enabled. Digestion was set to Trypsin/P and up to two missed cleavages were allowed. The FDR was set to 0.01 for peptides, proteins, and sites. Other parameters were used as pre-set in the software. The resulting protein groups were filtered to remove reverse (decoy), only identified by site, and potential contaminant protein groups.

### Data analysis and visualization

Further data analysis was performed in R using the following packages: tidyverse (Wickham et al. 2019) and sjmisc (Lüdecke 2018) for data manipulation; ggrepel (Slowikowski 2024) and patchwork (Pedersen 2024, v.1.3.0.9000) for data visualisation; org.Hs.eg.db and biomaRt (Durinck et al. 2005; Durinck et al. 2009) for human genome annotations; clusterProfiler (Xu et al. 2024), enrichplot (Yu 2018) for GO term analysis and visualisation; and knitr (Xie 2017) for markdown file formatting. UniProt, Enterez and Ensembl gene IDs were accessed from Bioconductor (Carlson 2017) (org.Hs.eg.db v.3.18.0.). GO term analysis was conducted using the Gene Ontology database (Ashburner et al. 2000), (Gene Ontology Consortium et al. 2023), (Thomas et al. 2022). Mitochondrial membership was defined as appearance in the Mitocarta dataset (Rath et al. 2021).

Proteomics data imputation was based on the idea of assigning the type of missingness from the Protti library (Quast et al. 2022) and adapted. First, we calculated the number of the missing datapoints for each condition and protein and then assigned the type of missingness as one of four types: 1) 3 NAs = all missing; 2) 2 NAs = missing not at random (MNAR); 3) 1 NA = missing at random (MAR); 4) 0 NAs = complete cases. We then compared the type of missingness for both conditions and each protein and if they were not identical (both MAR or both complete), assigned a combined type of missingness as follows: 1) If all were missing in one group and all complete in the second, we assigned MNAR; 2) If all were complete in one group but MNAR in the second, we assigned it as MNAR; 3) In all other cases we dropped the protein from the dataset due to insufficient measurements. Lastly, we imputed the proteins MNAR from a down-shifted normal distribution and the protein values MAR using the ludovic method from the protti package (Quast et al. 2022). The proteomics data was then normalized using the EigenMS method (Karpievitch et al. 2009) and proteins with less than 2 unique peptides removed.

The Lipidomics dataset was normalized using total sum scaling, i. e. dividing each value by its column sum and then multiplying by the average of all column sums. For both datasets, intensity values were log2-transformed to achieve normal distribution and fold changes were calculated as the difference between average log2 intensities. The significance of the calculated fold changes was determined using a two-sided Student’s t-test with a p-value cutoff of 0.05. The significance of an increase or decrease in fold change of the groups of lipid samples, categorized by their head groups or characteristics of the fatty acid chains, was assessed using a one-sample Student’s t-test and a Wilcoxon signed-rank test with a p-value cutoff of 0.05.

### Analysis of published proximity labeling data

We generated specificity plots and correlation plots based on the data from Supplementary Table 4A of Cell Metabolism, Vol.32 (3), Antonicka H, Lin Z-Y, Janer A, Aaltonen MJ, Weraarpachai W, Gingras A-C, Shoubridge EA, A High-Density Human Mitochondrial Proximity Interaction Network, 479–497.e9, Copyright Elsevier (2020) (Antonicka et al. 2020) using the web tool ProHits-viz (Knight et al. 2017). In the specificity analysis, we adjusted the readout column by the length of the prey sequence (X-axis). We colored based on the following conditions: 1) purple dots - the gene appeared in both mitochondria and peroxisomes; 2) orange dots - the gene appeared in mitochondria; 3) blue dots - the gene appeared in peroxisomes; 4) yellow dots - other genes. We labeled genes that appeared in both organelles or only in peroxisomes.

To generate the correlation plots, we also used the ProHits-viz web tool to calculate the correlation between bait and bait, bait and prey, and prey and prey. We used the adjustment by prey sequence length and for the bait-bait correlation plot we kept only rows correlating highly with OCIAD1 (MTCH1_iso2-C-ter, MARCH5-N-ter, MARCH5-C-ter, SLC25A46-N-ter, MTCH2-N-ter, MTCH1_iso2-N-ter, MTCH1_iso1-N-ter, RHOT2-N-ter, FIS1-N-ter, OCIAD1-N-ter, OCIAD1-C-ter, MAVS-N-ter, FKBP8-N-ter, PTPN1-N-ter), to take a closer look at the correlation value in bait-bait. The bait-prey plot was generated using the filter by OCIAD1 C-term and OCIAD1 N-term and an abundance cutoff from 50 to 250.

## Acknowledgements

The authors would like to thank Dr. Mattia Morandi, Dr. Klaudia Maruszczak, and Dr. Michał Wasilewski for stimulating discussions and insights. The authors thank the proteomics core facility of IMol Polish Academy of Sciences led by Dr. Remigiusz Serwa for the important contribution to this work.

---

**Figure S1.**
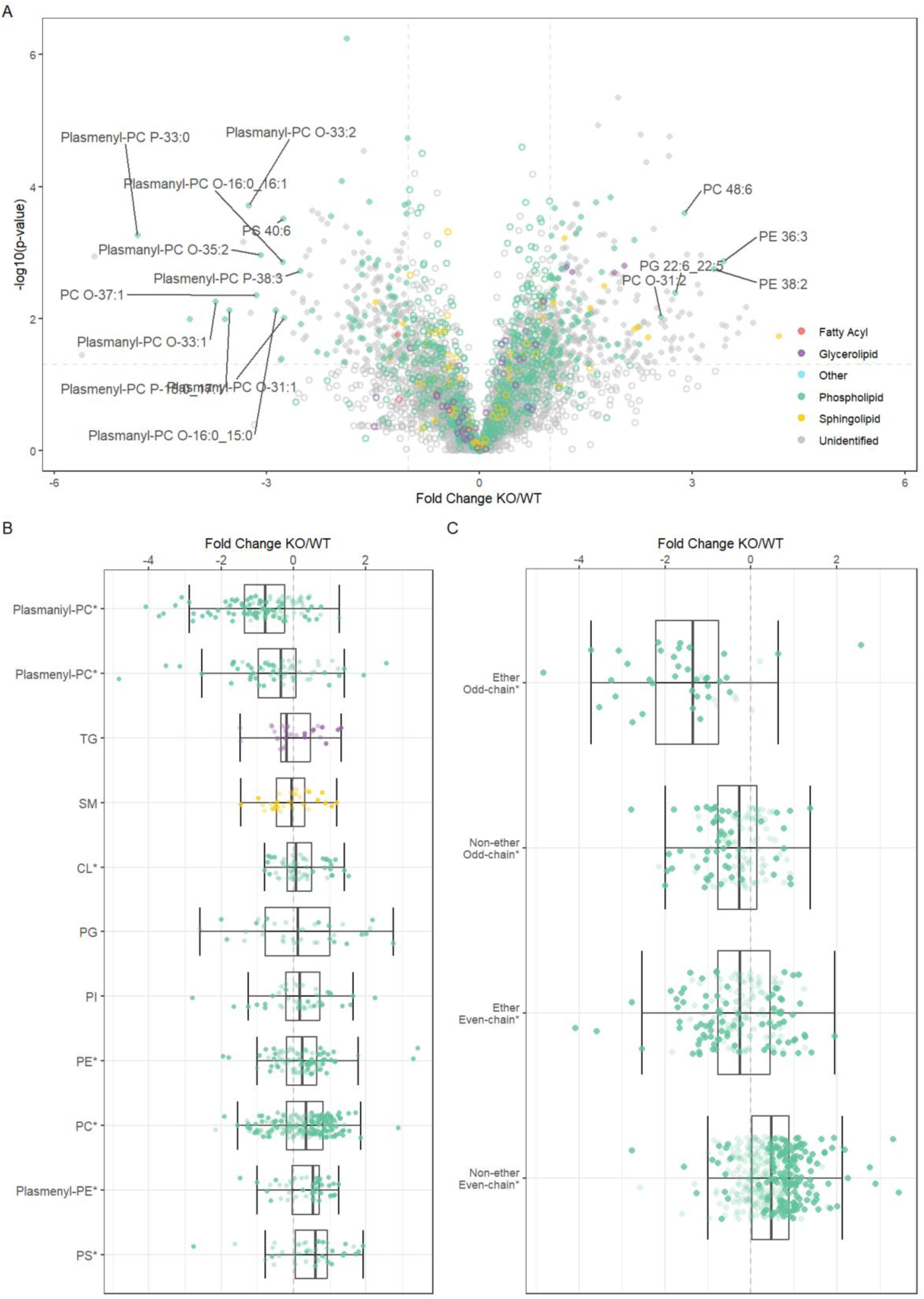
Lipidomics of OCIAD1 KO whole cells. Corresponding to Figure 1.

**Figure S2.**
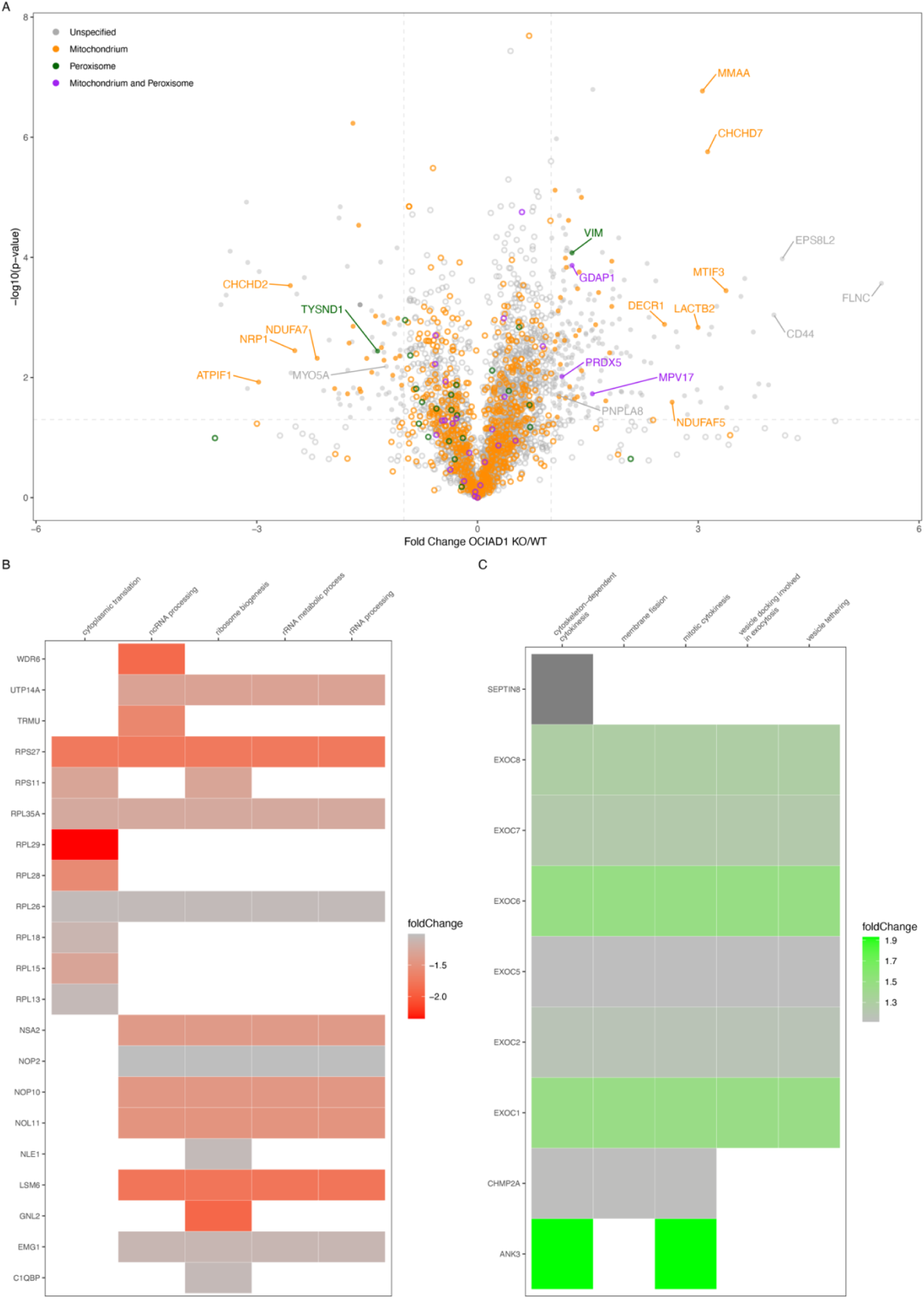
Proteomics of OCIAD1 KO whole cells. Corresponding to Figures 2 and 3. Proteomics of OCIAD1 KO whole cells compared to WT represented as volcano plot, colored by proteins of mitochondria (orange), peroxisome (green) or dually localized (purple) **(A)**. Heatmap of the top 5 enriched GO terms of proteins significantly decreased **(B)** and increased **(C)** upon OCIAD1 KO.

